# vep-rs: high-throughput Rust variant annotation with population-scale concordance to Ensembl VEP

**DOI:** 10.64898/2026.09.22.753614

**Authors:** Matthew Porter, Robert Borkowski

## Abstract

**Summary:** Ensembl VEP is the de facto reference for variant consequence annotation, but its Perl implementation limits throughput, and the open reimplementation benchmarked here is validated on a set too small to bound its disagreement with it. We present vep-rs, a Rust reimplementation of Ensembl VEP release 115.2, audited over 260,638,766 VEP consequence tuples, each a (location, allele, feature) key carrying a Sequence Ontology term set, from six variant datasets (ClinVar, gnomAD chr21, and 1000 Genomes chr21 across GRCh37 and GRCh38). On SNPs and indels vep-rs emits exactly VEP’s tuple count on all six and attains F1 ≥ 0.999974, differing from VEP on 3,169 tuples, at most one in 38,000 per dataset; with two documented VEP defect shapes set aside, F1 rounds to 1.000000 on all six. Per consequence class, 29 of the 30 Sequence Ontology terms carrying at least 1,000 tuples exceed F1 0.99, and the lowest, 0.540 on start_retained_variant, is entirely a VEP defect. On structural variants, F1 is 0.975395 (GRCh37) and 0.909998 (GRCh38). Timed on 20 independent machines per engine and architecture, vep-rs is faster than VEP by 101×-284× on ARM (geometric mean 176×) and 78.3×-192× on x86 (geometric mean 135×). Five divergence classes are documented, four of them defects in VEP release 115 that vep-rs does not reproduce: a consequence call that contradicts itself, a splice-region term dropped from a variant that lies in a splice region, annotation against a chromosome the variant does not lie on, and output that depends on input batch composition. A pipeline keyed on VEP consequence terms can therefore adopt vep-rs against a measured, per-term bound, shed VEP’s contradictory and batch-dependent calls, and annotate population-scale inputs at 78.3 to 284 times VEP’s throughput, and per-file structural-variant sets at 22.5 to 52.9 times.

**Availability and implementation:** vep-rs, its harness, both comparators, and every per-machine wall time are released under Apache-2.0 at https://github.com/natera-open-source/vep-rs (release v0.1.0, archived at https://doi.org/10.5281/zenodo.22837897).

**Supplementary information:** Supplementary data are available at *Bioinformatics* online.

## 1. Introduction

Sequencing cost has fallen faster than computing cost, so the secondary-analysis pipelines that turn reads into variant calls now hold a growing share of the cost and turnaround time of clinical sequencing (Berger and Yu, 2023). Interpretation workflows convert variant calls into transcript-level consequences before a filter or a curator acts on them, so the annotation vocabulary constrains the filters and reports built on it. Ensembl VEP (McLaren et al., 2016), hereafter VEP, is the de facto standard for that step and ships with the Ensembl reference releases. Its throughput is also a bottleneck at cohort scale: VEP annotates 10.96M gnomAD chromosome 21 records in 65.9 min with sixteen parallel workers on a 32-vCPU ARM host (Fig. 1b), and chromosome 21 is about 1.5% of the human genome.

**Figure 1.**
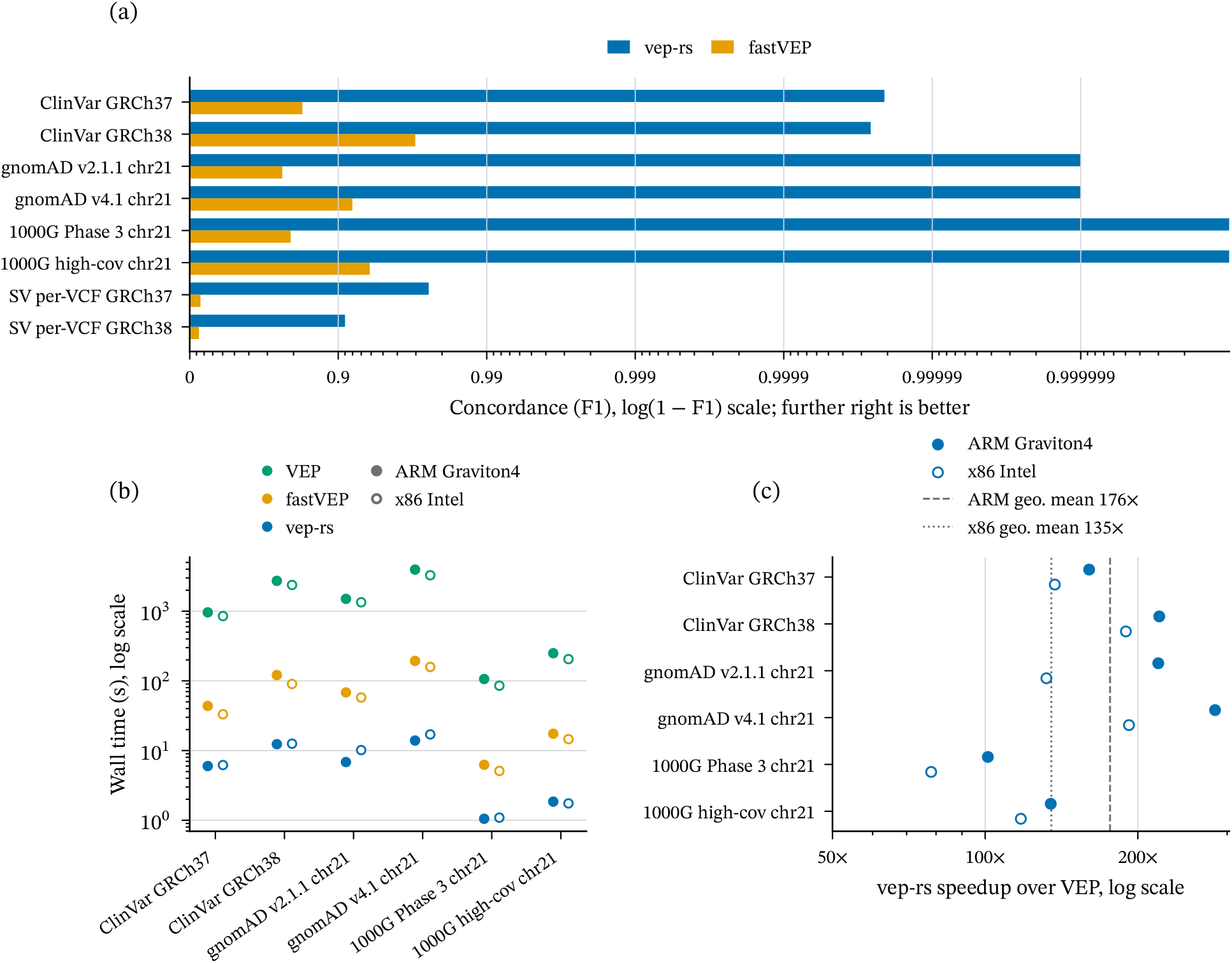
Concordance and throughput against Ensembl VEP release 115.2. (a) Raw F1 of vep-rs and fastVEP per variant dataset, the six SNP and indel datasets and the two per-assembly structural-variant sets, scored against the same VEP output on the tuple key of Section 2, on an axis logarithmic in 1 − F1; a longer bar is better, and a vep-rs bar reaching the right-hand limit is a dataset at F1 1.000000. (b) Wall time per engine on the six SNP and indel datasets on ARM Graviton4 and x86 Intel, median of 20 independent machine measurements after a discarded warmup, log scale; filled marks ARM and open marks x86. (c) Speedup of vep-rs over VEP on the same six datasets on both architectures, log scale, with reference lines at the ARM (dashed) and x86 (dotted) geometric means. Exact values are in Supplementary Tables S2, S4 and S6. Data: `manuscript/data/f1_observations.csv, f1_observations_fastvep.csv, wall_ times.csv`.

Faster annotators exist, but none of the four considered here is a drop-in replacement. SnpEff (Cingolani et al., 2012) and ANNOVAR (Wang et al., 2010) apply their own annotation rules, and ANNOVAR its own vocabulary, so a pipeline whose filters are written against VEP’s calls would have to revalidate them before adopting either. Nirvana (Stromberg et al., 2017) and fastVEP (Huang, 2026) target VEP’s vocabulary directly. Nirvana’s repository was archived in April 2026 under the PolyForm Strict licence, noncommercial use only, and its successor, Illumina Connected Annotations, is proprietary, so neither is open software a commercial clinical laboratory can adopt and Nirvana is not benchmarked here. fastVEP is open and is benchmarked throughout, but its published validation covers 2,340 transcript-allele pairs drawn from a 173-variant example file, four orders of magnitude fewer than the 33,483,881 tuples one ClinVar GRCh37 pass produces. A corpus of that size cannot resolve a rate finer than one in 2,340; on SNPs and indels vep-rs differs from VEP on 3,169 of its 260,638,766 tuples, one in 38,000 at worst (ClinVar GRCh38) and none of the 3,603,259 on 1000 Genomes Phase 3 chromosome 21.

vep-rs reimplements VEP in Rust for compiled performance, memory safety, and low-overhead parallelism. It emits VEP’s consequence vocabulary in VEP’s output formats, enabling a pipeline keyed on those terms to substitute it without rewriting its filters. This work supplies the missing bound over 260,638,766 VEP consequence tuples, reports that bound for each consequence class separately, and documents five classes of the residual disagreement, four of them defects in VEP release 115.

## 2. Methods

### Corpora

Six variant datasets span the genome-wide NCBI ClinVar releases of 2026-03-02 (GRCh37) and 2026-03-21 (GRCh38) (Landrum et al., 2018) and four chromosome 21 population subsets: gnomAD v2.1.1 and v4.1 genomes (Karczewski et al., 2020; Chen et al., 2024) and 1000 Genomes (1000G) Phase 3 (The 1000 Genomes Project Consortium, 2015) and high-coverage (Byrska-Bishop et al., 2022), each on the assembly its source publishes; the two 1000G call sets also carry symbolic structural alleles. Supplementary Table S1 gives every dataset’s input record count and consequence-tuple count. The structural-variant (SV) corpus is separate: 16 VCFs per assembly, 13 of them generated for this work to cover 41 VCF variant categories, three of those files being small-variant controls, plus chromosome 21 subsets of ClinVar, gnomAD-SV (Collins et al., 2020; Chen et al., 2024), and the 1000 Genomes structural-variant call sets (Sudmant et al., 2015; Byrska-Bishop et al., 2022). Supplementary S1.10 describes their construction; each file is annotated in its own run.

### Hardware and timing

All three engines, VEP release 115.2, fastVEP v0.3.0 and vep-rs, ran on AWS EC2 r8gd.8xlarge (ARM Graviton4) and r8id.8xlarge (x86 Intel), spec-matched at 32 vCPU and 256 GiB with the dataset on local NVMe. Per architecture and engine, 20 separately provisioned machines each ran one discarded warmup and one timed measurement per dataset (Supplementary S1). VEP ran inside its official container image and the two Rust engines natively. Every wall time is one engine annotating one dataset, the two structural-variant sets as 16 per-file invocations timed together (Supplementary S4.5), each a container start for VEP.

### Concordance metric

Each output line yields a key of location, allele, feature, feature type, and consequence set, the term sets sorted and deduplicated, ensuring equivalent orderings compare equal. Writing V for the tuples of VEP’s reference output, generated once per input (Supplementary S1.9), and *E* for the scored engine’s,

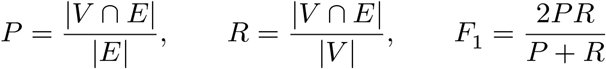

F1 is used because the engines need not emit the same number of tuples, one existing per variant, allele and transcript overlapping or within 5 kb, or one intergenic tuple where none does. An intersection over the scored engine’s count rewards emitting fewer rows, and over VEP’s count rewards emitting more. Where the two counts coincide, precision, recall, and F1 coincide too. The structural-variant comparator builds the same key after normalising contig names and breakend alleles; its F1 is the same metric computed over a different and much smaller corpus. Adjusted F1 recomputes *P* and *R* after removing from *V* and *E* the tuples the five documented divergence classes of Section 4.2 mask, four classes in full and one in part, four of them VEP defects and one a representation difference; it answers what concordance would be with those tuples set aside, and it is reported alongside raw F1 (Supplementary S2). Per-class F1 conditions the same three quantities on one Sequence Ontology term *t*, with *V*_*t*_ and *E*_*t*_ the tuples on each side whose term set contains *t*, a tuple counting for *t* only when its whole term set matches. F1 on this key scores the transcript rows each engine emits and the consequence terms on them. It does not certify VEP’s other output fields, and no VEP plugin enters any measurement here (Supplementary S1.3).

## 3. Implementation

vep-rs is nine released Rust crates in one Cargo workspace, seven of them in three layers and two cache tools beside them. A foundation crate carries the canonical types and a Consequence enum carrying all 41 terms of VEP’s Sequence Ontology vocabulary, of which vep-rs emits the 34 transcript-level and intergenic terms. Above it sit consequence calculation, the input parsers and output formatters, indexed FASTA access, eleven built-in plugins, and a C FFI loader for third-party plugins. The top layer is the binary and its pipeline runner, which batches 5,000 variants through rayon (Matsakis and Stone, 2025), emits VEP’s default, tab, VCF and JSON output formats in input order, field for field as VEP release 115.2 writes them for the same input under default flags, apart from installation-specific header values and the rows on which the two engines’ consequence calls differ, checked on a 1,754-record corpus for the tab, VCF and JSON formats and on all six datasets for the default format, and materialises transcript data per chromosome on first query. The two cache tools build transcript caches from Ensembl GFF3 and FASTA, re-lay an existing JSON cache and convert plugin annotation files to a memory-mapped store. Every vep-rs F1 here was measured on a Perl-derived cache, to hold transcript data fixed (Supplementary S1).

## 4. Results

### 4.1 Concordance

vep-rs reproduces VEP release 115.2 to at least F1 0.999974 on all six variant datasets of Supplementary Table S1, across 260,638,766 VEP tuples on the key Section 2 defines (Fig. 1a; Supplementary Table S2); it emits exactly the rows VEP emits, so all 3,169 disagreements are consequence-set differences on shared rows (Supplementary S2). ClinVar GRCh38 is the lowest of the six; the four chromosome 21 population sets reach 0.999999 or better, and the two 1000 Genomes sets 1.000000. Setting aside the two VEP defect shapes of Section 4.2 raises that bound to 1.000000 on all six.

When F1 is computed on each Sequence Ontology term separately, pooled across the six datasets, the aggregate conceals a per-class gap. Of the 30 terms carrying at least 1,000 tuples, 29 exceed F1 0.99 and one does not: start_retained_variant at 0.540 (Supplementary Fig. S1 and Table S3). It sits below the threshold because of the start co-emission defect of Section 4.2: all 1,270 VEP-only start_retained_variant tuples are pairs on which VEP emits start_lost and start_retained_variant together and vep-rs the same term set less the one the allele does not support, start_lost on the 1,269 sequence-variant tuples whose edited sequence still begins with the start codon and start_retained_variant on the one symbolic deletion that removes it; the mask excludes every one, so the term’s adjusted F1 is exactly 1. The class is rare, 2,759 tuples against 260,638,766, and its 2,539 one-sided tuple incidences move the aggregate by less than one part in a hundred thousand. fastVEP’s F1 is exactly zero on nine of the terms VEP emits, five of them above the same 1,000-tuple threshold. It emits no tuple at all for four of the nine; on the rest none of its tuples match VEP’s, on three terms because the consequence set differs and on transcript_ablation and transcript_amplification because only the location or allele does (Supplementary S4.2).

On structural variants, scored per VCF against the same VEP release over the 16 VCFs per assembly, raw F1 is 0.975395 on GRCh37 and 0.909998 on GRCh38, and adjusted F1 0.998524 on GRCh37 and 0.998031 on GRCh38 (Supplementary Table S4). Per file, adjusted F1 is at or above 0.999 on 13 of the 16 GRCh37 files and 14 of the 16 GRCh38, and exactly 1.0 on 10 and 8 files respectively (Supplementary Table S5).

fastVEP, on the same inputs and scored against the same VEP output, spans 0.762022 to 0.969701 on the six SNP and indel datasets and 0.133436 to 0.153626 on the structural-variant sets (Fig. 1a), a separation from vep-rs of 0.030 to 0.238 in absolute F1 on the former. Two caveats bound that comparison; on SNPs and indels the two transcript sets differ in both directions. fastVEP’s Ensembl GFF3 carries every transcript in VEP’s cache plus, on the canonical contigs, the 1,122 (GRCh37) and 2,134 (GRCh38) readthrough and artifact transcripts VEP’s cache build excludes, and the tuples fastVEP emits against those account for at most 0.004 of the separation; but its loader never loads 79,681 GRCh37 and 1,118 GRCh38 transcripts the cache holds, 88% to 96% of the VEP tuples it misses on the three GRCh37 datasets name one of them, and with those set aside its GRCh37 raw F1 would rise by 0.143 to 0.157 (Supplementary S1.8). On structural variants it emits 38% fewer tuples on GRCh37 and 12% fewer on GRCh38, and only a fifth and a seventh of those match VEP’s (Supplementary S4.3 and S4.5).

### 4.2 Divergence between vep-rs and VEP

Five divergence classes are documented here as the classes adjusted F1 sets aside. Each was identified from the measured disagreements and admitted on a criterion that does not ask whether vep-rs is right: in four, VEP’s output contradicts itself or the input it was given, the VEP defects; in the fifth, two self-consistent readings of one record differ only in representation. They account for all but 11 of the disagreements on SNPs and indels, 3,158 of the 3,169 tuples VEP emits that vep-rs does not, and the mask excludes 3,154 of the 3,169, or 99.5%; every divergence outside the classes, and the 4 inside them the mask does not reach, is charged to vep-rs in both the raw and the adjusted column. vep-rs reproduces none of the four defects.

Two reach SNPs and indels. For each differing region of a variant VEP tests the overlapping introns in turn and assigns rather than accumulates the splice_region_variant verdict, so a later region or intron that does not qualify overwrites an earlier one that did, dropping the term from a variant that does lie in a splice region. VEP can also emit both start_lost and start_retained_variant for the same allele, which cannot both hold.

Two reach structural variants. VEP annotates a structural variant against transcripts on a chromosome the variant does not lie on, so a deletion is reported as intronic to, flanking, or truncating a transcript it cannot reach. It removes 22,481 VEP-side tuples on GRCh38 and 1,838 on GRCh37. Separately, VEP’s transcript selection above --max_sv_size is batch-dependent, because it fills the transcript cache in 1 Mb bins on demand from whichever variants share the input batch: one giant breakend record yields 0 annotation rows in a batch of one, 22 in a batch of 82, and 1,281 once the threshold is raised past its span. On the whole 4,921-record gnomAD-SV chr21 input the same record yields 649, 224 and 50 rows at --fork 1, 4 and 16. VEP’s contract is to skip a variant wider than --max_sv_size, so the mask’s vep-rs-side removal is vep-rs’s complete annotation of the gnomAD-SV records above 10 Mb. Three of the four are traced to their source subroutines in the Ensembl release 115 code in Supplementary S2; the cross-chromosome class is defined on VEP’s output alone and its mechanism is not established.

The fifth class is a representation difference in which neither VEP nor vep-rs is wrong. On <CNV:TR> tandem repeats VEP expands the symbolic allele to its literal repeat bases and re-runs point-variant predicates over them, while vep-rs annotates it symbolically, a gain as feature_elongation and a contraction as feature_ truncation. Neither contradicts itself, and a symmetric mask removes all 1,368 swap pairs on GRCh37 and all 2,644 on GRCh38. <NON_REF> divergences are excluded from neither denominator, the comparator defining no filter for them.

Whether fastVEP reproduces four of the five is not decidable on this evidence; the fifth, the start co-emission, it cannot reproduce, since it never emits start_retained_variant (Supplementary Table S3). fastVEP loads only part of the transcript set VEP read (Section 4.1) and emits no counterpart for 3% to 33% of VEP’s tuples on the six SNP and indel datasets and 87.5% on the two structural-variant sets (Supplementary S4.1), so a defect absent from its output cannot be distinguished from one it fixed.

On SNPs and indels each exclusion removes one tuple from each side, shrinking both denominators equally. On structural variants the two sides’ removals are unequal, and the larger one raises F1 further. Removing only the smaller side’s count from both denominators, a conservative reading of the same classes, gives 0.984827 on GRCh37 and 0.929475 on GRCh38. Raw F1 is therefore the comparable quantity across engines, and adjusted F1 is best read within one (Supplementary S2).

### 4.3 Throughput

We timed all three engines on spec-matched ARM Graviton4 and x86 Intel hosts, vep-rs in one campaign and the comparators’ SNP and indel cells in another, each of separately provisioned machines, each figure the median of 20 measurements after a discarded warmup (Section 2); every ratio against VEP on those six datasets carries a between-campaign component not separated from the engine difference, and the same holds for every ratio against fastVEP (Supplementary S3.2). Inputs are stripped of per-sample genotypes, which cannot change a consequence set; the cost of carrying them is measured separately (Supplementary S4.6).

vep-rs is faster than VEP on every dataset on both architectures, by 101× to 284× on ARM (geometric mean 176×) and 78.3× to 192× on x86 (geometric mean 135×), and faster than fastVEP on every one as well, by 5.96× to 13.9× and 4.68× to 9.28× (Fig. 1b, 1c; Supplementary Table S6). No wall-time distribution’s P95 exceeds its P5 by more than 1.124× (Supplementary S1.2). Peak resident memory for vep-rs spans 0.62 to 4.56 GiB (Supplementary S1.4); VEP runs as sixteen forked processes inside a container, so only its cgroup peak is measurable (Supplementary S1.4).

The speedup over VEP persists when the input is not batched into a single invocation: the structural-variant sets are annotated as 16 separate per-file invocations per assembly, most of them small, so per-invocation cost weighs far more heavily, and vep-rs is 22.5× to 52.9× faster than VEP there on both architectures (Supplementary Table S7).

The speedup over VEP varies 2.81-fold across datasets on ARM, from 101× to 284×, and the speedup over fastVEP 2.33-fold. Each GRCh38 dataset carries 2.73× to 9.98× more consequence tuples than its GRCh37 counterpart and takes vep-rs 1.60× to 2.06× longer, so vep-rs’s tuple throughput rises by 1.33× to 5.94× on each of the six GRCh38 measurements across both architectures; VEP’s falls only on the ClinVar pair, from 34,800 to 33,600 tuples per second on ARM, and rises by 1.20× to 4.09× on the other two. vep-rs annotates that pair on ARM in 6.00 s and 12.35 s, at 5.59M and 7.41M tuples per second.

## 5. Conclusion

vep-rs reproduces Ensembl VEP release 115.2’s consequence calls at F1 ≥ 0.999974 on SNPs and indels, a bound that holds across 260,638,766 tuples on all six datasets and rises to 1.000000 once two documented VEP defect shapes are set aside, and at 0.975395 and 0.909998 on the GRCh37 and GRCh38 structuralvariant sets, while annotating the six SNP and indel datasets 78.3× to 284× faster than VEP and the two structural-variant sets 22.5× to 52.9× faster. Per consequence class, 29 of the thirty terms carrying at least a thousand tuples exceed F1 0.99, and the remaining one reaches exactly 1 once the documented VEP defect behind it is excluded. Pipelines keyed on VEP consequence terms therefore gain two orders of magnitude of throughput in geometric mean against a measured, per-term bound on their disagreement. Four of the five documented divergence classes are defects in VEP release 115 that vep-rs does not reproduce, so adopting it also removes VEP’s self-contradictory and batch-dependent calls from the annotation.

The residual divergence outside the five documented classes is 11 SNP and indel tuple pairs and the per-file structural-variant remainder of Supplementary Table S5, localised but not explained. vep-rs ships eleven built-in plugins, but no F1 reported here exercises any of them, so their agreement with VEP’s own implementations is unmeasured. Bounding them against the same reference at the same population scale is a natural next step, and the harness and comparators released here are what it would need.

## Acknowledgements

Anthropic Claude models were used, under author direction, in writing the vep-rs source, its measurement harness, its comparators, and the figure renderers.

## Author Contributions

M.P.: software, methodology, investigation, validation, data curation, visualization, writing (original draft). R.B.: software, methodology.

## Data availability

Apache-2.0 at https://github.com/natera-open-source/vep-rs, release v0.1.0, archived at https://doi.org/10.5281/zenodo.22837897, with the harness, both comparators, the figure renderers, every per-machine wall time, and the tuple counts every published F1 re-derives from. The engines’ annotation outputs on the benchmark datasets are not released: VEP’s reference outputs are retained as archival objects and the two Rust engines’ outputs behind the published figures are not retained; every published aggregate is in the released CSVs (Supplementary S3.3).

