## Supplementary Information for "vep-rs: high-throughput Rust variant annotation with population-scale concordance to Ensembl VEP"

Supplement to "vep-rs: high-throughput Rust variant annotation with population-scale concordance to Ensembl VEP" (Porter and Borkowski). Section references of the form "Section 4.1" point into the main text; "S1" through "S4" are sections of this document. "VEP" throughout means Ensembl VEP, the reference implementation; the release measured here is 115.2. This document carries the methods, the provenance, and the supporting tables for the concordance and wall-time figures the main text reports.

### S1 Methods

#### S1.1 Hardware and instance provisioning

A cell is one engine on one variant dataset on one architecture, the unit every wall-time figure is reported at; there are 56 of them rather than 48 because the two multi-sample datasets of S4.6 are separate datasets from the sites-only pair they share a source with. A sweep is one measurement campaign, identified by a dated sweep id, and this work composes two of them (S3.1).

All three engines ran on separately provisioned machines, each taking a single timed measurement, spanning two hardware-matched AWS EC2 instance types: r8gd.8xlarge (ARM Graviton4, 32 vCPU, 256 GiB RAM) and r8id.8xlarge (x86 Intel, 32 vCPU, 256 GiB RAM). The benchmark dataset was staged on each host's local NVMe instance store, preventing network-block-storage variance from entering the timed annotation. Matching vCPU count, memory, and storage class across the two instance families leaves the processor architecture and its per-architecture builds as the intended difference between them. VEP's measured container peak on the four sites-only inputs probed is at most 17.7 GiB (S1.4).

On each architecture, 20 fresh independent clones per engine, each a separately provisioned machine, booted from a stock Amazon Linux 2023 base image (`ami-0a2a049c945b84826` on arm64, `ami-029a761f237195c2c` on x86_64), self-bootstrapped every dependency and the full dataset from object storage, and self-terminated on result upload.

The official VEP container image `ensemblorg/ensembl-vep:release_115.2` compiles htslib (Bonfield et al., 2021) and its Perl binding from source and publishes native `linux/arm64` and `linux/amd64` manifests. VEP was timed from that image's native manifest on each architecture, without emulation, alongside the two Rust engines, which are built per architecture.

#### S1.2 Timing protocol and stability gate

Every wall-time cell is the median of N = 20 measurements per architecture: one timed run from each of 20 independent clones, each performing one warmup run that is discarded and then one timed measurement, both reading the dataset from local NVMe. Dispersion is carried in `manuscript/data/wall_times.csv` as an aggregate row of min, P5, P25, P75, P95, max, and the P95/P5 ratio on all 56 cells, alongside the 20 per-machine observations each of those rows recomputes from; the P95/P5 ratio is also tabulated beside each cell in Tables S6 to S8. Ratios of medians and geometric means are computed from the unrounded medians and printed to three significant figures.

All 56 cells' percentiles are computed under one estimator, the Weibull plotting position (R type 6; Hyndman and Fan, 1996), from the 20 per-machine observations each cell contributes.

Every cell clears the 1.3× P95/P5 stability gate. The highest observed ratio is 1.124 (vep-rs, x86, 1000G Phase 3 chr21); no cell was re-run on account of the gate. Recomputing every cell under the six continuous quantile definitions, R types 4 to 9, gives a maximum between 1.098 and 1.124, so no definition brings any cell near the gate.

Per-machine dataset parity is enforced for all three engines by an inventory digest that must match across all 20 clones of an engine on a given architecture; every clone of every cell rolled one inventory digest per engine, identical within and across both architectures (S3.1). The harness records each engine invocation's exit status alongside its wall clock rather than aborting the clone on a non-zero exit, so a failed run enters the data as an explicit failure instead of vanishing. Every timed cell exited zero, and every VEP cell also passed the harness's output check, which fails a run whose output holds fewer annotation rows than the input has records.

#### S1.3 Engine invocation

On every timed cell, vep-rs and VEP both run `--fork 16 --buffer_size 5000`; the 32-vCPU host leaves headroom above 16 forks. fastVEP does not accept a `--fork` flag and parallelises internally, set to 16 threads. The structural-variant cells of S4.5 run that same setting on each of their per-VCF invocations.

All engines consume canonical-filtered inputs restricted to Ensembl-style contigs (1-22, X, Y, MT), sites-only on every dataset of Tables S6 and S7: the six headline datasets are stripped of genotypes as below where their source carries them, and the one structural-variant input that carried a genotype matrix, the GRCh38 1000 Genomes file, is cut to its eight fixed columns (S1.10). VEP keys on position, allele, feature, and feature type, and genotypes do not affect the consequence set, so per-sample genotypes are stripped with `bcftools view -G` (Danecek et al., 2021) to keep the cross-engine comparison fair. The genotype-carrying cost is measured separately in S4.6.

No plugin was enabled on any engine in any measured run. Every F1 and wall time in this work is a core-annotation figure, and plugin behaviour enters none of them.

Beside every SNP and indel F1 the comparator scores per-column agreement on the rows both engines emit: the eight non-key columns (`Uploaded_variation`, `Gene`, `cDNA_position`, `CDS_position`, `Protein_position`, `Amino_acids`, `Codons`, `Existing_variation`) and the Extra keys `IMPACT`, `DISTANCE`, `STRAND` and `FLAGS`, with `OverlapBP` and `OverlapPC` on the two 1000G sets, agree on every shared row of all six datasets (`docs/concordance-provenance/2026-09-20-n2.json`, `field_agreement`).

#### S1.4 Peak resident memory

Each machine records peak resident memory alongside the wall clock, so the peak-memory figures here and in Section 4.3 are per-cell medians over that cell's own 20 measurements, on the same machines that produced its wall time, for vep-rs and fastVEP. Section 4.3 reports the vep-rs span only. Peak resident memory spans 0.62 to 4.56 GiB for vep-rs and 0.61 to 3.55 GiB for fastVEP across all cells; vep-rs is lighter on four of its ten cells on ARM (gnomAD v4.1, 1000G high-coverage in both conditions and the GRCh38 structural-variant set) and on those four plus 1000G Phase 3 with genotypes on x86, and fastVEP on the rest. Each engine's memory figures come from the same campaign as its wall times (S3.1). Values are binary gibibytes, converted from the kibibytes `/usr/bin/time -v` reports as `Maximum resident set size`. The x86 medians agree with the ARM medians to within 9.93% on every vep-rs cell and 0.27% on every fastVEP cell, the largest absolute cross-architecture difference being 0.0697 GiB on the vep-rs 1000G Phase 3 chr21 with genotypes cell.

VEP is the exception, for two reasons: it executes inside a container, so the harness observes only the `docker run` client process and its peak resident set is not captured by `/usr/bin/time -v` on that process, and it runs as sixteen forked processes, whose largest single resident set would not be comparable with a threaded engine's in any case. Its peak was therefore measured separately from the container cgroup's `memory.peak` at `--fork 16` on the timed ARM host class, giving 5,331 MiB, or 5.2 GiB, on the 1000G high-coverage GRCh38 cell and at most 18,124 MiB, or 17.7 GiB, across the four sites-only inputs probed (ClinVar GRCh38, gnomAD v4.1, 1000G Phase 3 and 1000G high-coverage). `memory.peak` charges file-cache pages to the container; the anonymous peak on the same four probes is 3,314, 5,827, 659 and 4,257 MiB, at most 5.7 GiB. On the same input carrying its 3,202 sample genotypes the peak is 25,323 MiB, or 24.7 GiB (S4.6). Those readings are single runs rather than medians of 20, outside both retained sweeps, and are the five rows of `manuscript/data/perl_bench.csv`. They bound the order of magnitude and are not comparable to the medians of Section 4.3, so peak memory is reported for the two engines the harness instruments directly and not for VEP.

#### S1.5 Tuple key construction and F1 computation

For each output line the harness constructs the five-tuple key Section 2 defines. Consequence sets are sorted and deduplicated before comparison, ensuring biologically equivalent term orderings are treated as identical. Tuples are deduplicated within each engine and intersected, by `sort -u` under `LC_ALL=C` and a streaming sort-merge for the SNP and indel datasets and by in-memory sets for the structural-variant files. Precision is the intersection over the scored engine's count, recall the intersection over the VEP count, and F1 is the harmonic mean 2PR/(P+R). Every F1 is an exact quotient of integer tuple counts and is printed to six decimals, except that a per-class value quoted in the text is rounded to three, since a class of a few thousand tuples does not resolve the finer places; Table S3 prints the six.

Implementation: `scripts/concordance/compare_vep_outputs.py`, `iter_vep_key_lines()` for key construction, `normalize_consequence_set()` for the sort and deduplication, and `prf()` for the precision, recall, and F1 formula. The structural-variant comparator `scripts/validation/compare_sv_concordance.py` constructs the same key with the same set normalisation, so the F1 in Table S4 is the same metric as the F1 in Table S2. In practice both engines emit consequence terms in Sequence Ontology rank order, so the normalisation guards against relying on that ordering rather than correcting any measured value.

#### S1.6 Per-dataset reporting and aggregates

The eight benchmark datasets (six SNP and indel, two per-assembly structural-variant) are not statistically independent. ClinVar appears in both the GRCh37 and GRCh38 cells; gnomAD v2.1.1 (GRCh37) and v4.1 (GRCh38) overlap in their underlying variant pool; 1000G Phase 3 (GRCh37) and high-coverage (GRCh38) overlap in samples. F1 is therefore reported per dataset and per assembly, and no arithmetic mean across datasets is reported, since that would weight overlapping datasets disproportionately.

The same non-independence applies to the wall-time speedups, which the main text does summarise across the six SNP and indel datasets by a geometric mean. That figure is a descriptive summary of these six cells and not an estimate of a population speedup over some wider workload. What the candidate estimators yield on the ARM cells of Table S6, every figure recomputed from the unrounded medians behind that table (S1.2):

| Statistic | vs VEP | vs fastVEP |
|---|---|---|
| Geometric mean | 176× | 9.07× |
| Arithmetic mean of ratios | 187× | 9.39× |
| Ratio of summed wall times | 226× | 10.7× |

On the x86 cells the geometric means are 135× against VEP and 6.55× against fastVEP; the main text prints the two against VEP.

Where a single aggregate is wanted, the tuple-weighted micro-F1 pools the tuples from all six datasets, then computes the intersection, both denominators, and F1 once on the pooled set. On the six SNP and indel datasets vep-rs reaches raw F1 0.999988, over 260,638,766 VEP tuples, 260,638,766 vep-rs tuples and a 260,635,597-tuple intersection; fastVEP reaches raw F1 0.920036 on the same six.

Every published F1 re-derives from committed integers rather than being a stored decimal: pool the `perl_tuples`, `rust_tuples`, and `intersection` columns of `manuscript/data/f1_observations.csv` for vep-rs and the corresponding columns of `manuscript/data/f1_observations_fastvep.csv` for fastVEP, subtract a dataset's excluded-pair count from both raw denominators for an adjusted SNP and indel value (S2), and read the two structural-variant rows' post-mask denominators from their own columns. Every published raw and adjusted F1 recomputes from those integers by that arithmetic. Table S5's per-file values re-derive the same way from the per-file counts in `manuscript/data/sv_concordance_by_category.csv`.

#### S1.7 Cache parity between the two engines

Both engines read the same transcript data: VEP its Storable cache, and vep-rs a JSON conversion of that cache produced offline by `scripts/data/storable_to_json.pl` and shipped as its runtime input; the GRCh37 conversion is of the cache-version-113 Storable, whose 310 differences from the 115 build S1.9 bounds. The converted caches carry 195,232 GRCh37 and 505,231 GRCh38 transcripts, each assembly's Ensembl gene set on the canonical contigs less the readthrough and artifact transcripts VEP's cache build skips (S1.8); on GRCh38 chromosome 21, where both caches were listed directly, the Storable cache holds the same 6,752 transcripts as the JSON cache (`manuscript/data/transcript_set_parity.csv`). Because the VEP path consumes the original Storable while vep-rs consumes the JSON, a field-coverage gap in the converter would be a systematic source of F1 bias. A converter audit over a seeded sample of N = 1000 GRCh37 transcript records drawn from a randomised chunk order, 903 of them on the canonical contigs the JSON cache carries (`scripts/concordance/cache_parity_audit.pl`; manifest `docs/concordance-provenance/2026-09-16-cache-parity-audit.json`), found every transcript field VEP's consequence pipeline consumes present on both sides but one, the sequence-edit list of the next paragraph: exon and intron coordinates, CDS bounds, peptide sequences and protein identifiers, biotype, gene metadata, and the variant-effect feature cache including its three-prime and five-prime UTR substructures. Fields differing by a documented type-level conversion, Perl objects becoming plain JSON objects and sequence objects becoming strings, are classified as converted-under-rule rather than as gaps.

The Storable cache's SIFT and PolyPhen prediction matrices are not carried into the JSON conversion, and no F1 in this work reads them. The per-transcript sequence edits (`seq_edits`) VEP applies when it rewrites a reference peptide are not carried either, but their effect is: the cached peptide is VEP's own translation with those edits applied, and vep-rs reads the edited residues from it.

#### S1.8 Comparability of the fastVEP cells

All three engines consume canonical Ensembl-style VCFs per cell: the F1 comparisons read pre-generated canonical files, staged on every clone and pinned by the dataset inventory digest (S1.2), whose UCSC `chr` prefixes were stripped at generation time by `scripts/concordance/prepare_benchmark_vcfs.py`, and the VEP wall-time runs read a copy the harness derives from the raw input by the same contig strip and genotype removal. The comparators key on the location string, so contig naming is normalised upstream of every engine; the structural-variant comparator additionally normalises contig names and breakend alleles at comparison time (Section 2). fastVEP's raw output is then passed through `scripts/concordance/fastvep_normalize.py` before comparison, which rewrites intergenic `Feature_type=Transcript` to `-`

so that both engines' rows key alike; the comparison fails if that normaliser is absent, rather than falling back to the un-normalised output. fastVEP's transcript caches were built from Ensembl GFF3 annotation, `Homo_sapiens.GRCh37.87.chr.gff3` and `Homo_sapiens.GRCh38.115.gff3`, the GRCh37 gene set being frozen at release 87. That normaliser is applied to the SNP and indel output only; for the structural-variant cells fastVEP's symbolic alleles are mapped to VEP's Sequence Ontology class names before comparison, and those cells are not like-for-like, for the reason S4.5 gives.

Compared by transcript identifier, each GFF3 carries every transcript of the matching VEP-derived cache, 195,232 on GRCh37 and 505,231 on GRCh38, and the cache holds none the GFF3 lacks. The GFF3 carries more: on the canonical contigs 1,122 (GRCh37) and 2,134 (GRCh38) transcripts, exactly the readthrough transcripts, and on GRCh38 the 19 of biotype `artifact`, that VEP's cache build skips (`ensembl-vep` release 115.2, `Bio/EnsEMBL/VEP/AnnotationSource/Database/Transcript.pm`, lines 266 to 270; verified against the Ensembl release 115 core databases), and on GRCh38 a further 2,285 on scaffolds outside the canonical contigs the inputs are restricted to. fastVEP annotates against those transcripts: of the tuples it emits that VEP does not, 5.8% (gnomAD v2.1.1) to 25.6% (ClinVar GRCh37) name one, and no tuple VEP emits does. Removing them from fastVEP's denominator would raise its raw F1 by 0.002271 to 0.003759, so of the 0.030 to 0.238 separation of Table S2 at most 0.004 is attributable to the source difference. What fastVEP loads from its GFF3 is a different matter: its loader (v0.3.0, `crates/fastvep-cache/src/gff.rs`) admits sixteen transcript-level GFF3 feature types, reads gene, exon and CDS records separately, and discards transcript records of every other type, so it never loads 79,681 of the GRCh37 cache's transcripts (`processed_transcript` 27,762, `aberrant_processed_transcript` 25,872, `lincRNA` 11,752, `processed_pseudogene` 11,058, `pseudogene` 3,212, `nc_primary_transcript` 25) and 1,118 of the GRCh38 cache's (`unconfirmed_transcript` 1,106, `processed_transcript` 12). Of the tuples VEP emits that fastVEP does not, 92.9% (ClinVar GRCh37), 87.7% (gnomAD v2.1.1) and 96.0% (1000G Phase 3) name one of those transcripts, against 1.4%, 0.8% and 0.6% on the three GRCh38 datasets. Removing those VEP tuples from VEP's count with the intersection fixed would raise fastVEP's raw F1 from 0.825597 to 0.968830, 0.762022 to 0.911212 and 0.790928 to 0.948071 on GRCh37 and by at most 0.0003 on GRCh38, so on GRCh37 most of fastVEP's separation from vep-rs is transcripts its loader never reads, and on GRCh38 almost none of it is; the remainder on both, beyond the at most 0.004 the source difference contributes, is fastVEP's own annotation. Data: `manuscript/data/transcript_set_parity.csv` (columns `cache_transcripts_fastvep_admitted`, `cache_transcripts_fastvep_unloaded`), `manuscript/data/transcript_set_parity/<assembly>_fastvep_unloaded.tsv`, `manuscript/data/fastvep_extra_tuples_by_transcript.csv` (column `vep_only_on_fastvep_unloaded_transcripts`).

#### S1.9 Regenerating the reference output

The figure and CSV pipeline consumes pre-computed F1 values. To regenerate them end to end: build or convert the transcript cache, canonicalise the inputs to Ensembl-style contigs, generate the VEP release 115.2 reference output per canonical input with `--fasta` parity at every call site, then run each engine and compare tuple-by-tuple with the two comparators of S1.5. The single measurement entrypoint is `scripts/concordance/run_clone_measurement.sh`, which runs every dataset for one engine against a populated data tree; `scripts/concordance/run_concordance.sh` is the development-time harness that runs a benchmark directory's VCFs through both engines locally. The SNP and indel reference output is a canonical VEP release 115.2 set at cache version 115, the ClinVar GRCh37 set at cache version 113; the two GRCh37 caches hold the same 195,232 transcripts, 194,922 of them identical in content and 310 differing in the stored peptide's first residue, in protein-domain features or, on 235 of them, in the SIFT and PolyPhen matrices no F1 reads (S1.7), and a ClinVar GRCh37 reference regenerated at version 115 reproduces the 113 reference output byte for byte, 33,483,881 tuple keys on each side (`manuscript/data/transcript_set_parity.csv`; `docs/concordance-provenance/2026-09-18-grch37-cache-113-vs-115.json`); the structural-variant reference output, the set `sv_per_vcf_r115.2_c115_tr_regen_20260920` on both assemblies, annotates each of the 16 VCFs per assembly independently at `--fork 4 --buffer_size 5000` (`docs/concordance-provenance/2026-09-09-sv-grch37-reference.json` records that invocation), a setting the reference depends on because VEP's transcript selection above `--max_sv_size` follows batch composition (S2). Per-consequence-class F1 (Table S3) is derived from the same archived artefacts with no engine re-run: per-term VEP totals stream from the canonical VEP output, per-term one-sided counts come from each engine's archived `report/discordant.tsv`, and the per-term intersection is the VEP total less the VEP-only count.

#### S1.10 Structural-variant corpus construction

The 13 synthetic structural-variant VCFs are generated for this work; there is no external corpus to cite. `scripts/validation/generate_sv_test_vcfs.py` emits them deterministically per assembly from one specification, making a file's variant count shared between GRCh37 and GRCh38 while its coordinates are assembly-specific. The 13 generator functions produce 41 VCF variant categories in total; three files (`01_snv`, `02_mnp_complex`, `03_small_indels`) are small-variant controls. Every record is placed on chromosome 21 except the 100 mate records of `09_breakends`'s inter-chromosomal pairs, which are placed on chromosomes 1 to 5 so that annotation across a breakend junction is exercised on both assemblies.

| File | Variants | Categories covered |
|---|---|---|
| `01_snv` | 1,000 | 1 |
| `02_mnp_complex` | 2,000 | 2 |
| `03_small_indels` | 2,000 | 2 |
| `04_large_explicit` | 400 | 2 |
| `05_symbolic_del_ins` | 2,000 | 2 |
| `06_symbolic_dup_inv` | 2,250 | 4 |
| `07_cnv_repeat` | 900 | 3 |
| `08_mobile_elements` | 800 | 5 |
| `09_breakends` | 1,400 | 7 |
| `10_special_alleles` | 1,000 | 4 |
| `11_multi_allelic` | 700 | 2 |
| `12_complex_imprecise` | 300 | 2 |
| `13_vcf45_features` | 650 | 5 |
| Total | 15,400 | 41 |

The three real-world files are chromosome 21 subsets of ClinVar release 2026-03-02 restricted to its structural `CLNVC` classes (Deletion, Duplication, Insertion, Inversion, copy number gain and copy number loss), of gnomAD-SV v2.1 on GRCh37 and v4.1 on GRCh38, and of the 1000 Genomes Phase 3 structural-variant release v8 (2013-05-02) on GRCh37 and the high-coverage structural-variant freeze V3 (2021-01-24) on GRCh38, filtered to canonical contigs under the same policy as the SNP and indel inputs (Ensembl-style contigs 1-22, X, Y, MT); the GRCh38 1000 Genomes file is cut to its eight fixed columns, so every structural-variant input is sites-only. Each of the 16 files per assembly is annotated in its own engine invocation, so no file's transcript-cache population can affect another's and the batch-composition class of S2 stays observable per file.

### S2 Adjusted-F1 exclusions

Raw F1 scores every tuple as a binary match. Adjusted F1 additionally excludes tuples matched by a documented divergence class. Three of the five classes exclude matched consequence-swap pairs, removing one tuple from the VEP and one from the vep-rs denominator; the other two, both structural-variant classes, exclude single one-sided tuples, the cross-chromosome class on the VEP side only. Every exclusion leaves the intersection unchanged, so an adjusted value can only rise, and on the structural-variant sets the two denominators lose different numbers of tuples; the aggregate counts are below. Four classes are fully masked and one in part, and those five are the whole of the mask. Four of the five are VEP defects on the criterion Section 4.2 states; the fifth is the `<CNV:TR>` representation difference of Section 4.2. Divergences no class excludes are charged to vep-rs in both the raw and the adjusted column and are not itemised as classes here; S4.2 breaks one term's residual down by shape. On the six SNP and indel datasets vep-rs emits exactly the rows VEP emits, so every one of the 3,169 divergences is a consequence-set difference on a row both engines carry (`manuscript/data/f1_observations.csv`, whose VEP-only and vep-rs-only counts coincide on every dataset, and the per-dataset pair count in `docs/concordance-provenance/2026-09-20-n2.json`).

The five classes, with the tuple count each accounts for per assembly, the two SNP and indel rows on ClinVar alone:

| Class | Mask status | GRCh37 | GRCh38 | Basis |
|---|---|---|---|---|
| `sv_nondeterministic_transcript_selection` | fully masked | 10,934 | 183,746 | exact |
| `sv_cross_chromosome_transcript_annotation` | fully masked | 1,838 | 22,481 | exact |
| `start_cooccurrence_swap` | fully masked | 261 | 928 | exact |
| `sv_cnv_tr_literal_expansion` | fully masked | 1,368 | 2,644 | exact |
| `splice_family_swap` | partially masked (440 of 440; 1,444 of 1,444; the class's 4 unmasked pairs are on gnomAD v2.1.1 chr21) | 440 | 1,444 | exact |

The rows must not be summed, because they do not share one basis. The transcript-selection and cross-chromosome counts are one-sided exclusions, the `<CNV:TR>` row counts symmetric pairs, one tuple from each side, the two SNP and indel rows are ClinVar swap-pair counts rather than structural-variant tuples, and the `splice_family_swap` row is a class size of which only the parenthesised fraction is excluded. The taxonomy CSV's `masked_pairs` column spans those tuple grains for the same reason and cannot be pooled into either denominator. Every row is measured at the published binary. Data: `manuscript/data/discordance_taxonomy.csv`.

The two SNP and indel defects are in `ensembl-variation` at release/115, commit `23c76f60` (github.com/Ensembl/ensembl-variation), which carries VEP's consequence-calling modules; the batch-dependent defect is in `ensembl-vep` at the same release, commit `2beada0d` (github.com/Ensembl/ensembl-vep), which carries its input and cache layer and the structural-variant input path on which the cross-chromosome class arises. The `splice_region` overwrite is `BaseTranscriptVariationAllele.pm`, `sub _intron_effects` at line 99, whose nested per-region and per-intron loops assign the verdict at line 215 rather than taking a logical disjunction, so a later region or boundary intron overwrites an earlier positive one; of that subroutine's fourteen writes it is the only one that assigns a return value, which is why the mechanism can drop exactly one term, `splice_region_variant`. The start-codon co-emission is `Utils/VariationEffect.pm`, `sub start_lost` at 851, `sub start_retained_variant` at 947, and `sub _` ins_del_start_altered at 1028, where a DNA-level retention check disagrees with a peptide-level loss check and VEP emits that mutually exclusive pair for one allele. The retention check is the one the edited sequence supports: the pair arises only when _ins_del_start_altered finds the edited 5′ UTR and coding sequence still beginning with the start codon, and start_lost then fires through the codon-window peptide translation, which does not see the upstream base that re-forms the codon. On 10:56424022-56424024 (deletion, ENST00000320301) VEP emits 5_prime_UTR_variant, start_lost and start_retained_variant for a deletion the UTR absorbs with the coding sequence unchanged, and vep-rs 5_prime_UTR_variant and start_retained_variant; on the one symbolic deletion in the class, which does remove the codon, start_lost is the supported term and vep-rs keeps it. The cross-chromosome class is defined on VEP's output alone, a named transcript that the shared transcript cache does not hold for the variant's chromosome, and arises on the structural-variant path that begins in VEP/Parser/VCF.pm:create_StructuralVariationFeatures; no line is established as its mechanism. The batch-dependent transcript selection is VEP/AnnotationSource.pm:get_all_features_by_InputBuffer, which populates the transcript cache in genomic bins of cache_region_size, 1 Mb by default, on demand from whichever variants share the input batch; the replay Section 4.2 cites is docs/concordance-provenance/2026-09-16-bnd-batch-replay.json.

The mask does not cover its reporting classes exactly. It under-excludes on the one partly masked class. The splice_region overwrite can drop exactly one term, so the mask excludes only the pairs that mechanism explains: of the 1,888 splice-family swap pairs it accounts for all 440 on ClinVar GRCh37 and all 1,444 on GRCh38 and for none of the 4 on gnomAD v2.1.1 chr21, on which vep-rs adds splice_donor_5th_base_variant rather than splice_region_variant; those 4 stay in the denominators and depress vep-rs's F1.

The batch-dependent class is masked on a proxy, because batch dependence cannot be seen in a single run's output. A tuple is excluded when the other engine annotated the same variant but not against this transcript, vep-rs named at least as many transcripts for that variant as VEP did (strictly more for a vep-rs-side tuple), and the variant's span exceeds VEP's 10 Mb --max_sv_size default. A transcript-set difference meeting all three can still potentially have another cause.

The cross-chromosome class reaches both assemblies, removing 22,481 VEP-side tuples on GRCh38 and 1,838 on GRCh37, which is nearly all of the gap between the two assemblies' VEP-side exclusion totals; the larger gap between their vep-rs-side totals, 12,302 against 186,390, belongs almost entirely to the batch-dependent class. Whether a tuple belongs to the class is objectively checkable, the shared transcript cache either holding the named transcript for the variant's chromosome or not, but the mechanism producing it is not established here. The GRCh38 gnomAD-SV input carries chr-prefixed CHR2 values against its stripped CHROM while the GRCh37 input's CHR2 values match its CHROM, and the class removes 1,838 tuples on GRCh37 all the same, so the contig-naming mismatch does not produce it. One record is listed in docs/concordance-provenance/2026-09-19-sv-cross-chromosome-example.json, from the archived reference output: a 25 kb deletion at 21:23699593-23724683 for which VEP names 122 transcripts, 120 of them on chromosome 22.

On SNPs and indels, the mask removes a specific divergence shape. A swap pair is excluded only when the term set vep-rs adds contains splice_region_variant, lies within {splice_region_variant, intron_variant}, and VEP holds no term vep-rs lacks; a start-codon co-emission pair, an inherent contradiction, is excluded when vep-rs's term set is VEP's less one of the two start terms, which is the shape of every one of the 1,270 co-emission pairs measured (S4.2).

| Dataset | Excluded from both denominators |
| --- | --- |
| ClinVar GRCh37 | 701 |

| Dataset | Excluded from both denominators |
| --- | --- |
| ClinVar GRCh38 | 2,372 |
| gnomAD v2.1.1 chr21 | 3 |
| gnomAD v4.1 chr21 | 77 |
| 1000G Phase 3 chr21 | 0 |
| 1000G high-cov | 1 |

Subtracting a dataset's count from both raw denominators of `manuscript/data/f1_observations.csv` reproduces its published adjusted F1; the 1000G Phase 3 cell needs no mask at all, so its raw and adjusted F1 coincide. Summed across the six datasets the mask removes 3,154 of the 3,169 tuples VEP emits that vep-rs does not, 99.5%. The five classes contain 3,158 of the 3,169; the 11 outside them are eight tuples at one start codon on chromosome 20 (two SNVs on four transcripts), an SNV in a mitochondrial start codon on both ClinVar sets and one 6 kb ClinVar GRCh38 deletion, and the 4 in-class pairs the mask does not reach are the gnomAD v2.1.1 splice-family pairs above; all 15 are charged to vep-rs in both columns.

On the structural-variant classes the per-side totals diverge, because the two engines disagree by different amounts in each direction. On GRCh37 the adjusted metric removes 12,302 vep-rs-side and 3,206 VEP-side tuples across all masked classes combined; on GRCh38, 186,390 and 25,125. Each of those four figures is the difference between the raw and adjusted denominators in `manuscript/data/f1_observations.csv`; the two VEP-side figures are the cross-chromosome row plus the `<CNV:TR>` row of the table above, and the two vep-rs-side figures the transcript-selection row plus the `<CNV:TR>` row. With the intersection unchanged, $F1 = 2I/(|V| + |E|)$ depends only on the total number of tuples removed, so a larger removal raises it further whichever side it falls on. The final column below removes the smaller side's count from both denominators, a conservative reading of the same exclusion sets that sets aside the 9,096 (GRCh37) and 161,265 (GRCh38) further tuples the one-sided removals take out.

| Dataset | Raw | Published adjusted | Adjusted under a symmetric exclusion |
| --- | --- | --- | --- |
| SV GRCh37 | 0.975395 | 0.998524 | 0.984827 |
| SV GRCh38 | 0.909998 | 0.998031 | 0.929475 |

The vep-rs-side removal is 3.84 times the VEP-side one on GRCh37 and 7.42 times on GRCh38, so on GRCh38 the vep-rs-side removal supplies most of the movement from 0.909998 to 0.998031. A symmetric reading of the same classes reaches 0.929475, and the remaining 0.068556 comes from the 161,265 further tuples the one-sided removals take out. Neither column is the corrected value the classes would produce if the divergences were fixed rather than excluded; they bound what the exclusion itself contributes.

Three of the five mask predicates match the shape of a consequence-set difference rather than a property of the variant and cannot distinguish a term either engine adds correctly from one it adds wrongly. The other two do not read the consequence set: the batch-dependent proxy keys on transcript-set size and a span gate above VEP's `--max_sv_size` default, and the cross-chromosome predicate keys on whether the shared transcript cache holds the named transcript for the variant's chromosome. This is why the raw columns are the comparable pair across engines and the adjusted columns are best read within an engine.

### S3 Provenance

#### S3.1 Pinned artefacts

| Item | Value |
|---|---|
| Wall-time and concordance sweeps | Two campaigns. `sweep-20260920T012411Z`: every vep-rs cell (all wall time, all concordance, both Table S8 conditions) and the two structural-variant cells of fastVEP and VEP, wall time and fastVEP concordance. `sweep-20260918T224520Z`: the SNP and indel cells of fastVEP and VEP (wall time, fastVEP concordance and both Table S8 fastVEP conditions), measured on the same instance types; the two campaigns share the SNP and indel inputs and reference outputs, and every structural-variant cell in this work comes from the later one. Every campaign carries a dataset-inventory digest per engine. |
| vep-rs binary digest | `23a294cf` (ARM) / `48b2a959` (x86), built natively per architecture from the crate sources of release v0.1.0 (Zenodo, https://doi.org/10.5281/zenodo.22837897) at two target-cpu codegens, every clone reporting the same digest; the release assets are separate builds verified by their own `SHA256SUMS` |
| vep-rs build inputs | `cargo build --release --locked`, rustc 1.97.0 (2d8144b78 2026-07-07), the release's `Cargo.lock` sha256 `d3961d1e`… |
| fastVEP release | `v0.3.0`, binary digest `6d83c5bc` (ARM) / `e123fbc0` (x86) |
| fastVEP output normaliser | `scripts/concordance/fastvep_normalize.py`, md5 2698a8a7 |
| VEP engine | `ensemblorg/ensembl-vep:release_115.2`, image digest `sha256:ff3c7e20`… |
| Input policy | `universal_ensembl_style`, canonical-contig filter active |
| Reference output (SNP and indel) | canonical VEP release 115.2, cache version 115 (ClinVar GRCh37 at cache version 113, which a reference regenerated at 115 reproduces byte for byte; S1.9) |
| Reference output (SV) | per-VCF VEP release 115.2, 16 VCFs per assembly: `sv_per_vcf_r115.2_c115_tr_regen_20260920` on both assemblies, generated against the full-genome Ensembl caches |
| Dataset inventory digest | `sweep-20260920T012411Z`: vep-rs `15c55e95`…, fastVEP `41797ac1`…, VEP `cde7eee3`…; `sweep-20260918T224520Z`: fastVEP `6e287b12`…, VEP `be81ccea`…; each identical across all 40 clones of its engine in its campaign on both architectures, the two campaigns' digests differing on the structural-variant inputs and reference (S1.10); within a campaign the engines' digests differ because each engine's inventory covers its own transcript cache (the JSON conversion, the GFF3-built cache and the Storable cache respectively), and fastVEP's additionally covers the `drivers/` prefix it stages |
| Clones per cell | 20 fresh clones per engine per architecture, one timed measurement each |
| Base images | `ami-0a2a049c945b84826` (arm64), `ami-029a761f237195c2c` (x86_64), stock Amazon Linux 2023 |

Concordance is deterministic per binary and architecture-independent, and was verified on the full cohort. On every measured dataset, every clone that scored it produced identical raw F1 and tuple counts to six decimal places: all 20 vep-rs clones per architecture on every dataset and all 20 fastVEP clones per architecture on the six SNP and indel datasets and the two structural-variant sets. Adjusted F1 is identical across the same clones on the six SNP and indel datasets; on the two structural-variant sets it is read from 2 further clones per architecture per engine and is identical across them (`sv_scoring_note` in the concordance record, `fastvep_cross_clone_determinism` in the wall-time record). The released form of that check is the per-clone manifest in `docs/concordance-provenance/2026-09-20-n20-parallel-clones.json`, one entry per clone per engine and architecture, the vep-rs and fastVEP entries pinned to their binary digest and dataset inventory digest and the VEP entries to the campaign's image digest and dataset inventory digest, with the vep-rs verdict in the companion record `2026-09-20-n2.json` and the fastVEP verdict, per dataset with the number of clones reporting, in the wall-time record's `fastvep_cross_clone_determinism` field; the per-clone concordance reports the verdicts were computed from are withheld archival objects (S3.3).

#### S3.2 Table columns spanning more than one sweep

Every cross-engine ratio on the six SNP and indel datasets divides medians measured in separate campaigns rather than side by side: vep-rs was timed in one sweep and both comparators' SNP and indel cells in another, on the same instance types against the same dataset inventory (S3.1), so a between-campaign component is present in each of those ratios and is not separated from the engine difference. The two structural-variant cells of every engine were measured in the later sweep, so Table S7's ratios carry no between-campaign component, and every ratio still divides medians from separate machines. Every vep-rs cell, both Table S8 conditions included, comes from one campaign; each comparator's cells come from two, its structural-variant pair from the later one. `manuscript/data/wall_times.csv` records the source sweep of every cell.

#### S3.3 Released and withheld artefacts

Released in the repository: the measurement harness (`scripts/concordance/`, including the single measurement entrypoint `run_clone_measurement.sh`; the clone provisioning scripts that stage the data and upload results are not released), both comparators (`scripts/concordance/compare_vep_outputs.py`, `scripts/validation/compare_sv_concordance.py`) and their test suites, the input pointers and canonical-VCF preparation scripts, the structural-variant validation VCFs and their generator, every per-cell aggregate and per-clone observation behind each table (`manuscript/data/*.csv`, 1,176 wall-time rows covering 20 timed measurements for all 56 cells, the per-dataset tuple counts behind every aggregate F1 and the post-mask denominators on the two vep-rs structural-variant rows, the per-file tuple counts behind Table S5 for both engines, and the 800 container start-up timings behind S4.5), and the figure renderers. Figures regenerate end to end from the released CSVs via `python3 manuscript/figures/generate_figures.py --all`, and a digest manifest of the six renderer outputs committed under `manuscript/figures/` is written to `manuscript/figures/MANIFEST.txt`. The released artefacts are additionally deposited at Zenodo (https://doi.org/10.5281/zenodo.22837897) as the archive of release v0.1.0, so every repository-relative path cited here resolves inside the deposit unchanged; that DOI is the citation route for the data CSVs.

Not in the repository: VEP's reference annotation outputs and each clone's per-dataset comparison report, retained as archival objects; the vep-rs and fastVEP clones upload their comparison reports and timings, not their annotation outputs. Those objects, and the VEP campaign's per-clone timing files, are the per-clone provenance behind each released aggregate; every median, percentile ratio and adjusted-F1 denominator in this work recomputes from the released CSVs without them. They are held in institutional object storage rather than a public archive and are not distributed; the aggregates every published figure and table is computed from are in the released CSVs, so no number in this work depends on access to them.

#### S3.4 Use of language models

Anthropic Claude models were used, under author direction, in writing the vep-rs source, its measurement harness, its comparators, and the figure renderers. Every table value in this work is read from the released CSVs and both figures are rendered from them by the script named in S3.3, and the authors verified each published value against them.

### S4 Supplementary tables and figure

#### S4.0 Dataset inventory

**Table S1.** The six SNP and indel variant datasets and the two per-assembly structural-variant sets, with the input record count each canonical VCF carries and the consequence-tuple count VEP produces from it. Input records are counted on the canonical-contig-filtered VCF each engine consumes, sites-only, so they are the record counts the wall times of Tables S6 and S7 are measured against. VEP tuples are the distinct tuple keys in VEP's output (S1.5), summed over the 16 files of each structural-variant set (S4.5), which every F1 in Tables S2 and S4 is scored against.

| Dataset | Assembly | Source release | Input records | VEP tuples |
|---|---|---|---|---|
| ClinVar | GRCh37 | 2026-03-02 | 4,388,172 | 33,483,881 |
| ClinVar | GRCh38 | 2026-03-21 | 4,398,127 | 91,552,813 |
| gnomAD v2.1.1 chr21 | GRCh37 | v2.1.1 | 3,483,000 | 11,075,422 |
| gnomAD v4.1 chr21 | GRCh38 | v4.1 | 10,958,898 | 110,503,269 |
| 1000G Phase 3 chr21 | GRCh37 | Phase 3 | 1,105,538 | 3,603,259 |
| 1000G high-coverage | GRCh38 | high-cov | 1,002,753 | 10,420,122 |
| SV per-VCF | GRCh37 | ClinVar 2026-03-02 (CLNVC structural classes); gnomAD-SV v2.1; 1000G Phase 3 SV v8 2013-05-02; synthetic corpus | 23,992 | 329,864 |
| SV per-VCF | GRCh38 | ClinVar 2026-03-02 (CLNVC structural classes); gnomAD-SV v4.1; 1000G high-coverage SV freeze V3 2021-01-24; synthetic corpus | 53,212 | 1,117,037 |

Data: `manuscript/data/f1_observations.csv` for the tuple counts, `manuscript/data/dataset_inventory.csv` for the record counts.

#### S4.1 Consequence-tuple concordance

**Table S2.** Consequence-tuple concordance against VEP release 115.2, the six SNP and indel variant datasets, both engines. Both engines read the same input VCFs and are scored against the same VEP output, but their transcript data come from different sources: vep-rs reads a conversion of the cache VEP reads, while fastVEP builds its own from Ensembl GFF3, which carries every transcript that cache holds plus the readthrough and artifact transcripts VEP's cache build excludes, and whose loader never reads 79,681 GRCh37 and 1,118 GRCh38 of the cache's transcripts (S1.8).

| Engine | Dataset | Assembly | VEP tuples | Raw F1 | Adj F1 |
|---|---|---|---|---|---|
| vep-rs | ClinVar | GRCh37 | 33,483,881 | 0.999979 | 1.000000 |
| vep-rs | ClinVar | GRCh38 | 91,552,813 | 0.999974 | 1.000000 |
| vep-rs | gnomAD v2.1.1 chr21 | GRCh37 | 11,075,422 | 0.999999 | 1.000000 |
| vep-rs | gnomAD v4.1 chr21 | GRCh38 | 110,503,269 | 0.999999 | 1.000000 |
| vep-rs | 1000G Phase 3 chr21 | GRCh37 | 3,603,259 | 1.000000 | 1.000000 |
| vep-rs | 1000G high-cov | GRCh38 | 10,420,122 | 1.000000 | 1.000000 |
| fastVEP | ClinVar | GRCh37 | 33,483,881 | 0.825597 | 0.825605 |
| fastVEP | ClinVar | GRCh38 | 91,552,813 | 0.969701 | 0.969712 |
| fastVEP | gnomAD v2.1.1 chr21 | GRCh37 | 11,075,422 | 0.762022 | 0.762022 |
| fastVEP | gnomAD v4.1 chr21 | GRCh38 | 110,503,269 | 0.919675 | 0.919675 |
| fastVEP | 1000G Phase 3 chr21 | GRCh37 | 3,603,259 | 0.790928 | 0.790928 |
| fastVEP | 1000G high-cov | GRCh38 | 10,420,122 | 0.938604 | 0.938604 |

The separation between the two engines is 0.030 to 0.238 in absolute F1. fastVEP emits no counterpart for 2.7% (ClinVar GRCh38) to 32.9% (gnomAD v2.1.1 chr21) of the tuples VEP emits on the six SNP and indel datasets, and for 87.5% on each of the two structural-variant sets of Table S4, one minus its recall, from the `perl_tuples` and `intersection` columns of `manuscript/data/f1_observations_fastvep.csv`. The mask covers 99.5% of the tuples VEP emits that vep-rs does not and 0.0055% of those it emits that fastVEP does not, which is why S2 reads the adjusted columns within an engine only.

#### S4.2 Per-consequence-class concordance

Per-class F1 conditions precision and recall on one Sequence Ontology term, as Section 2 defines; for a tuple in the intersection the two term sets are identical by construction, because `consequence_set` is one of the five key columns. A term's per-class value is therefore not a partition of the aggregate and the column cannot be pooled back into it; term sets overlap, so one tuple contributes to several classes.

Every value re-derives from committed integers, as the aggregates do, by the derivation S1.9 describes.

The lowest per-class value is explained in full. Every one of the 1,270 VEP-only `start_retained_variant` tuples is a start co-emission pair, VEP emitting `start_lost` and `start_retained_variant` together where vep-rs emits the same term set less the start term the allele does not support (S2): on the 1,269 sequence-variant tuples vep-rs keeps `start_retained_variant`, so the term sits on both sides of the pair and the pair still differs, and on the one symbolic deletion it keeps `start_lost`; the `start_cooccurrence_swap` class of S2 excludes all of them, so the term's raw F1 of 0.540 is entirely that VEP defect and its adjusted F1 is exactly 1, and the 1,269 vep-rs-only `start_retained_variant` tuples are the vep-rs sides of those same pairs. The pairs also count against `start_lost`, 1,270 on the VEP side and one on the vep-rs side, because `consequence_set` is a key column, and account for all but 18 of its one-sided tuples; its raw F1 is 0.992 and its adjusted F1, with those 18 open, rounds to 1.000 (Table S3). Data: `manuscript/data/start_retained_shapes.csv`.

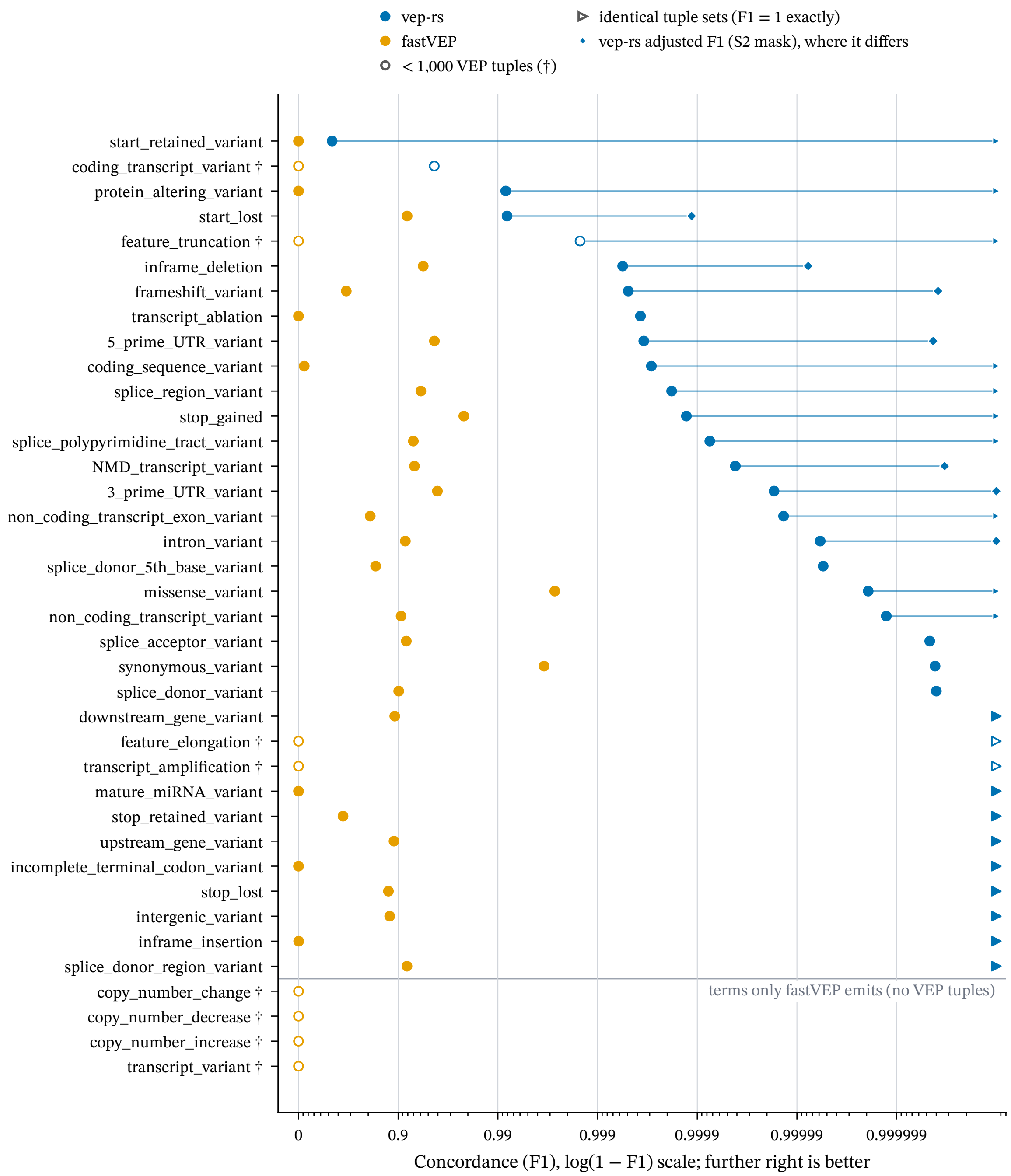


**Figure S1.** Concordance, F1, per Sequence Ontology consequence term against VEP release 115.2, both engines, pooled over the six SNP and indel variant datasets, on an axis logarithmic in $1 - \text{F1}$; a mark further right is better. Every term of Table S3 is drawn. The upper 34 rows are the terms VEP emits, ordered worst first on vep-rs raw F1, one mark per engine on raw F1 and, for vep-rs, a second smaller mark on adjusted F1 (S2) where it differs from raw; the four rows below the rule are terms only fastVEP emits. An open mark, matched by a dagger in the label, is a term below the 1,000-tuple threshold of Section 4.1. A right-pointing mark at the right-hand limit is a term whose two tuple sets are identical, F1 exactly 1, 11 of the 34 for vep-rs on raw F1 and 21 on adjusted (three further raw terms and eight adjusted values print 1.000000 at six decimals while differing by one to ten tuples); the axis floor is 1e-7. Table S3 gives every value with its tuple count.

**Table S3.** Per-consequence-class F1 for both engines, on the basis of Figure S1. Terms are ordered by VEP tuple count descending; † marks a term below the 1,000-tuple threshold of Section 4.1. Each value is the exact quotient of the integer counts in the data file, printed to six decimals; the text rounds a per-class value to three (S1.5). The vep-rs adjusted column recomputes each term's F1 after removing from both sides the pairs the S2 mask excludes; fastVEP's adjusted values are not tabulated, the mask reaching 0.0055% of its one-sided tuples (S4.1). The last four terms carry no VEP tuples at all: fastVEP emits them and VEP does not. Nine further classes read 0.000000 for fastVEP, for two different reasons. It emits no tuple at all for four of them. On the remaining five it does emit tuples, and none of them match: on three terms its consequence set differs from VEP's on every tuple, and on `transcript_ablation` and `transcript_amplification` the set matches on all but two tuples while the location or allele does not (`manuscript/data/fastvep_zero_f1_shapes.csv`). On `feature_elongation`, for example, it pairs the term with the Sequence Ontology copy-number term `copy_number_increase`, which release 115 does not emit, so its count equals its `copy_number_increase` count in each dataset, 174 and 1,774.

| Term | VEP tuples | vep-rs F1 | vep-rs adj F1 | fastVEP F1 |
|---|---|---|---|---|
| `intron_variant` | 128,705,338 | 0.999994 | 1.000000 | 0.915221 |
| `non_coding_transcript_variant` | 70,902,816 | 0.999999 | 1.000000 | 0.906487 |
| `missense_variant` | 35,783,494 | 0.999998 | 1.000000 | 0.997313 |
| `downstream_gene_variant` | 30,146,836 | 1.000000 | 1.000000 | 0.891712 |
| `upstream_gene_variant` | 25,250,015 | 1.000000 | 1.000000 | 0.889699 |
| `synonymous_variant` | 12,157,381 | 1.000000 | 1.000000 | 0.996560 |
| `NMD_transcript_variant` | 12,126,342 | 0.999959 | 1.000000 | 0.931323 |
| `non_coding_transcript_exon_variant` | 10,061,610 | 0.999986 | 1.000000 | 0.809229 |
| `3_prime_UTR_variant` | 5,629,571 | 0.999983 | 1.000000 | 0.959569 |
| `splice_region_variant` | 5,257,276 | 0.999819 | 1.000000 | 0.940773 |
| `intergenic_variant` | 4,927,601 | 1.000000 | 1.000000 | 0.878366 |
| `splice_polypyrimidine_tract_variant` | 4,916,495 | 0.999925 | 1.000000 | 0.929536 |
| `frameshift_variant` | 2,603,915 | 0.999508 | 1.000000 | 0.668088 |
| `stop_gained` | 1,494,776 | 0.999872 | 1.000000 | 0.978029 |
| `splice_donor_variant` | 1,251,188 | 1.000000 | 1.000000 | 0.901156 |
| `5_prime_UTR_variant` | 1,161,850 | 0.999656 | 1.000000 | 0.956686 |
| `splice_acceptor_variant` | 1,074,252 | 1.000000 | 1.000000 | 0.917095 |
| `splice_donor_region_variant` | 759,812 | 1.000000 | 1.000000 | 0.918404 |
| `splice_donor_5th_base_variant` | 458,373 | 0.999995 | 0.999995 | 0.831695 |
| `inframe_deletion` | 389,223 | 0.999440 | 0.999992 | 0.943917 |
| `inframe_insertion` | 155,083 | 1.000000 | 1.000000 | 0.004066 |
| `coding_sequence_variant` | 138,690 | 0.999712 | 1.000000 | 0.123917 |
| `start_lost` | 80,277 | 0.991907 | 0.999886 | 0.918652 |
| `stop_lost` | 28,706 | 1.000000 | 1.000000 | 0.874647 |
| `protein_altering_variant` | 20,718 | 0.991650 | 1.000000 | 0.000000 |
| `stop_retained_variant` | 20,310 | 1.000000 | 1.000000 | 0.642341 |
| `incomplete_terminal_codon_variant` | 3,990 | 1.000000 | 1.000000 | 0.000000 |
| `start_retained_variant` | 2,759 | 0.539786 | 1.000000 | 0.000000 |
| `transcript_ablation` | 1,346 | 0.999629 | 0.999629 | 0.000000 |
| `mature_miRNA_variant` | 1,111 | 1.000000 | 1.000000 | 0.000000 |
| `feature_truncation` † | 667 | 0.998501 | 1.000000 | 0.000000 |
| `transcript_amplification` † | 505 | 1.000000 | 1.000000 | 0.000000 |
| `feature_elongation` † | 208 | 1.000000 | 1.000000 | 0.000000 |
| `coding_transcript_variant` † | 12 | 0.956522 | 0.956522 | 0.000000 |
| `copy_number_change` † | 0 | — | — | 0.000000 |

| Term | VEP tuples | vep-rs F1 | vep-rs adj F1 | fastVEP F1 |
|---|---|---|---|---|
| copy_number_decrease † | 0 | — | — | 0.000000 |
| copy_number_increase † | 0 | — | — | 0.000000 |
| transcript_variant † | 0 | — | — | 0.000000 |

Data: manuscript/data/f1_by_consequence_class.csv for the pooled values printed here, and manuscript/data/f1_by_consequence_class_by_dataset.csv for the per-(dataset, term) detail the pooling sums.

#### S4.3 Structural-variant concordance

**Table S4.** Structural-variant concordance against per-VCF VEP release 115.2 reference output. Both engines are scored against the same 16 per-assembly VEP outputs, on the same VEP-side denominators, under the same exclusion classes (Section 4.2, S2). Two classes match on fastVEP: the cross-chromosome class, which is defined on VEP's own output and so removes the same VEP tuples for fastVEP as for vep-rs (1,838 on GRCh37 and 22,481 on GRCh38), and the <CNV:TR> class, on 87 and 165 symmetric pairs (excluded_categories in manuscript/data/f1_observations_fastvep.csv). The fastVEP separation from VEP here is only in part one of annotation volume: 41,166 of the 206,062 tuples it emits on GRCh37 match VEP's, and 140,071 of 982,409 on GRCh38 (S4.5).

| Engine | Dataset | Assembly | Variants | Raw F1 | Adj F1 |
|---|---|---|---|---|---|
| vep-rs | SV per-VCF | GRCh37 | 23,992 | 0.975395 | 0.998524 |
| vep-rs | SV per-VCF | GRCh38 | 53,212 | 0.909998 | 0.998031 |
| fastVEP | SV per-VCF | GRCh37 | 23,992 | 0.153626 | 0.154205 |
| fastVEP | SV per-VCF | GRCh38 | 53,212 | 0.133436 | 0.134902 |

**Table S5.** vep-rs structural-variant concordance per input file, against the per-VCF VEP release 115.2 reference output, each file on its own tuple counts, for the 13 synthetic and 3 real-world files of each assembly. Read each assembly's two columns against each other, not across assemblies. The 13 synthetic files are generated per assembly from one specification (S1.10), so a file's variant count is shared; the three real-world files carry their GRCh37 / GRCh38 counts.

| Input file | Variants | GRCh37 raw | GRCh37 adjusted | GRCh38 raw | GRCh38 adjusted |
|---|---|---|---|---|---|
| 01_snv | 1,000 | 1.000000 | 1.000000 | 1.000000 | 1.000000 |
| 02_mnp_complex | 2,000 | 1.000000 | 1.000000 | 0.999822 | 0.999822 |
| 03_small_indels | 2,000 | 1.000000 | 1.000000 | 1.000000 | 1.000000 |
| 04_large_explicit | 400 | 1.000000 | 1.000000 | 1.000000 | 1.000000 |
| 05_symbolic_del_ins | 2,000 | 1.000000 | 1.000000 | 0.999991 | 0.999991 |
| 06_symbolic_dup_inv | 2,250 | 1.000000 | 1.000000 | 1.000000 | 1.000000 |
| 07_cnv_repeat | 900 | 0.936322 | 1.000000 | 0.923333 | 0.999969 |
| 08_mobile_elements | 800 | 1.000000 | 1.000000 | 1.000000 | 1.000000 |
| 09_breakends | 1,400 | 0.997981 | 0.998969 | 0.998294 | 0.998992 |
| 10_special_alleles | 1,000 | 0.955571 | 0.955571 | 1.000000 | 1.000000 |
| 11_multi_allelic | 700 | 0.999533 | 0.999533 | 1.000000 | 1.000000 |
| 12_complex_imprecise | 300 | 1.000000 | 1.000000 | 0.999008 | 0.999008 |
| 13_vcf45_features | 650 | 0.999014 | 0.999014 | 0.999843 | 0.999843 |

| Input file | Variants | GRCh37 raw | GRCh37 adjusted | GRCh38 raw | GRCh38 adjusted |
|---|---|---|---|---|---|
| 1kg_sv_chr21 | 877 / 2,386 | 1.000000 | 1.000000 | 0.999762 | 0.999762 |
| clinvar_sv_chr21 | 2,794 / 2,796 | 0.999952 | 0.999952 | 1.000000 | 1.000000 |
| gnomad_sv_chr21 | 4,921 / 32,630 | 0.832168 | 0.990660 | 0.813283 | 0.995552 |

Thirteen of the 16 GRCh37 files and 14 of the 16 GRCh38 files reach adjusted F1 of 0.999 or better, and 10 and 8 respectively reach exactly 1.0. `gnomad_sv_chr21` carries nearly all of the transcript-selection and cross-chromosome classes of S2, so its two columns differ by 0.158 (GRCh37) and 0.182 (GRCh38); `07_cnv_repeat` carries the whole `<CNV:TR>` class, and its one divergence outside that mask, on GRCh38, is charged to vep-rs; every other file's two columns are equal or nearly so. Every value re-derives from the per-file VEP, vep-rs and intersecting tuple counts committed beside it, raw and adjusted, in `manuscript/data/sv_concordance_by_category.csv`, which carries fastVEP's per-file F1 and counts on the same basis; fastVEP is not tabulated per file here. Data: `manuscript/data/sv_concordance_by_category.csv`.

#### S4.4 Wall time

**Table S6.** Wall time on two spec-matched architectures, the six SNP and indel variant datasets, median of 20 independent clone measurements per cell per engine per architecture; the three P95/P5 columns give each cell's dispersion across its 20 measurements (S1.2). Structural-variant wall time is measured as 16 separate invocations and is reported separately in S4.5.

| Arch | Dataset | VEP (s) | fastVEP (s) | vep-rs (s) | Speedup vs VEP | Speedup vs fastVEP | VEP P95/P5 | fastVEP P95/P5 | vep-rs P95/P5 |
|---|---|---|---|---|---|---|---|---|---|
| ARM | ClinVar GRCh37 | 961.08 | 43.65 | 6.00 | 160× | 7.28× | 1.096 | 1.038 | 1.049 |
| ARM | ClinVar GRCh38 | 2,722.01 | 121.13 | 12.35 | 220× | 9.81× | 1.054 | 1.044 | 1.036 |
| ARM | gnomAD v2.1.1 chr21 | 1,500.91 | 68.24 | 6.84 | 219× | 9.98× | 1.044 | 1.018 | 1.012 |
| ARM | gnomAD v4.1 chr21 | 3,953.00 | 193.44 | 13.92 | 284× | 13.9× | 1.042 | 1.027 | 1.075 |
| ARM | 1000G Phase 3 chr21 | 106.29 | 6.26 | 1.05 | 101× | 5.96× | 1.048 | 1.047 | 1.110 |
| ARM | 1000G high-cov | 249.15 | 17.45 | 1.85 | 135× | 9.43× | 1.046 | 1.054 | 1.066 |
| x86 | ClinVar GRCh37 | 849.97 | 33.16 | 6.20 | 137× | 5.35× | 1.063 | 1.066 | 1.037 |
| x86 | ClinVar GRCh38 | 2,378.35 | 90.26 | 12.56 | 189× | 7.19× | 1.057 | 1.050 | 1.032 |
| x86 | gnomAD v2.1.1 chr21 | 1,341.62 | 57.69 | 10.16 | 132× | 5.68× | 1.050 | 1.035 | 1.057 |
| x86 | gnomAD v4.1 chr21 | 3,276.52 | 158.24 | 17.06 | 192× | 9.28× | 1.068 | 1.051 | 1.022 |

| Arch | Dataset | VEP (s) | fastVEP (s) | vep-rs (s) | Speedup vs VEP | Speedup vs fastVEP | VEP P95/P5 | fastVEP P95/P5 | vep-rs P95/P5 |
|---|---|---|---|---|---|---|---|---|---|
| x86 | 1000G Phase 3 chr21 | 85.31 | 5.10 | 1.09 | 78.3× | 4.68× | 1.087 | 1.060 | 1.124 |
| x86 | 1000G high-cov | 205.16 | 14.60 | 1.75 | 118× | 8.36× | 1.079 | 1.059 | 1.077 |

#### S4.5 Structural-variant wall time

Structural-variant wall time is excluded from the main text headline speedup claim and geometric mean. Each of the four cells annotates one assembly's full structural-variant set on one architecture as 16 separate per-VCF invocations: 13 small synthetic files covering 41 variant categories on chr21, with 100 breakend mates on chromosomes 1 to 5 (15,400 variants, S1.10), plus three real-world files (ClinVar, gnomAD, 1000G chr21). This section is also the per-invocation control Section 4.3 cites for the throughput claim.

**Table S7.** Structural-variant wall time, median of 20 independent clone measurements per cell per engine per architecture, with each cell's P95/P5 dispersion (S1.2). No vep-rs-over-fastVEP ratio is given, because the fastVEP column is not like-for-like (see below).

| Arch | Dataset | Assembly | VEP (s) | fastVEP (s) | vep-rs (s) | Speedup vs VEP | VEP P95/P5 | fastVEP P95/P5 | vep-rs P95/P5 |
|---|---|---|---|---|---|---|---|---|---|
| ARM | SV set | GRCh37 | 32.28 | 18.78 | 0.61 | 52.9× | 1.049 | 1.038 | 1.049 |
| ARM | SV set | GRCh38 | 64.92 | 72.18 | 2.81 | 23.1× | 1.072 | 1.043 | 1.082 |
| x86 | SV set | GRCh37 | 29.62 | 17.56 | 0.79 | 37.5× | 1.054 | 1.025 | 1.063 |
| x86 | SV set | GRCh38 | 59.29 | 68.65 | 2.63 | 22.5× | 1.044 | 1.018 | 1.061 |

These cells are not directly comparable with Table S6's. Each cell runs 16 process invocations over mostly small inputs, so per-invocation startup carries far more weight relative to annotation throughput than it does on a single population-scale pass. For VEP each invocation is a container start: a bare start of the pinned image, timed 20 times on each of the 40 VEP machines of the structural-variant campaign after their timed cells, has a median of 1.015 s on ARM and 0.950 s on x86 (`manuscript/data/container_start.csv`), so the 16 starts are about half of the 32.28 s GRCh37 cell on ARM and a quarter of the 64.92 s GRCh38 one, and the same shares on x86. The speedups span 22.5× to 52.9× over VEP across the four cells, below the 78.3× to 284× of the Table S6 corpus. On ARM, VEP itself takes 32.28 s and 64.92 s here, against 1,500.91 s and 3,953.00 s on the two gnomAD cells, so a 16-invocation corpus of small files bounds what any speedup can be measured against.

The fastVEP column is not like-for-like. Against the same per-VCF VEP reference output, fastVEP emits 206,062 consequence tuples on the GRCh37 set and 970,269 on GRCh38, against VEP's 329,830 and 1,103,635 and vep-rs's 339,588 and 1,267,435 (deduplicated cross-file union counts, a tuple key occurring in more than one of an assembly's 16 files counted once; the file-summed denominators behind Table S4 are 329,864 and 1,117,037 on the VEP side and 206,062 and 982,409 on fastVEP's, both bases columns of `manuscript/data/f1_observations_fastvep.csv`, the vep-rs union counts in the concordance record of S3.1). Doing 38% and 12% less tuple-level work than VEP on the same inputs, at correspondingly low concordance, means its wall time cannot be read as a speedup ratio against engines that annotate the full set.

Both engines of Table S4 are scored on the file-summed basis under one exclusion set. The cross-chromosome class is the one exclusion that treats them differently, as its assertion holds for one engine and not the other. That assertion is that the scored engine cannot name a transcript the reference cache lacks for the variant's chromosome. It is true of vep-rs, whose per-chromosome index is built from that cache, and not of an engine carrying its own annotation release, so on fastVEP the shape is counted, not excluded, and stays in both raw and adjusted F1, at 242 tuples on GRCh37 and 3,638 on GRCh38.

#### S4.6 Multi-sample sample-scaling

The headline throughput comparison is measured on sites-only inputs. Under the invocation used here VEP assigns consequences per position, allele and feature, so per-sample genotypes never enter the consequence calculation or the tab output; every cell of Tables S6 and S7 runs a sites-only input (S1.3). Stripping genotypes before the timed annotation is therefore the fair basis for cross-engine comparison. VEP was not timed in the genotype-carrying condition in its campaign, which is why Table S8 omits it. A single probe run of the same `--fork 16` invocation on the 3,202-sample 1000G high-coverage input completed on the timed ARM host class in 2,681 s, 10.8 times the 249.15 s sites-only median of Table S6, writing the same 10,420,122 annotation rows at a container peak of 24.7 GiB (S1.4; `manuscript/data/perl_bench.csv`).

**Table S8.** Multi-sample sample-scaling: median wall time in seconds, N = 20 independent clone measurements, for the two 1000G variant datasets, sites-only against per-sample genotypes retained, with each genotype-carrying cell's P95/P5 dispersion (S1.2; the sites-only cells' are in Table S6). The speedup column is the vep-rs-over-fastVEP speedup in the genotype-carrying condition. VEP is not tabulated (see above).

| Dataset (samples) | Assembly | Arch | vep-rs sites (s) | vep-rs +GT (s) | fastVEP sites (s) | fastVEP +GT (s) | Speedup vs fastVEP (+GT) | vep-rs +GT P95/P5 | fastVEP +GT P95/P5 |
|---|---|---|---|---|---|---|---|---|---|
| 1000G Phase 3 (2,504) | GRCh37 | ARM | 1.05 | 1.14 | 6.26 | 165.33 | 145× | 1.062 | 1.042 |
| 1000G Phase 3 (2,504) | GRCh37 | x86 | 1.09 | 1.43 | 5.10 | 127.56 | 89.2× | 1.109 | 1.106 |
| 1000G high-coverage (3,202) | GRCh38 | ARM | 1.85 | 2.08 | 17.45 | 201.59 | 96.9× | 1.039 | 1.046 |
| 1000G high-coverage (3,202) | GRCh38 | x86 | 1.75 | 2.38 | 14.60 | 154.60 | 65.0× | 1.081 | 1.056 |

The genotype-carrying cost is a real deployment consideration for pipelines that annotate cohort VCFs in place, and the gap between the two engines widens once genotypes are carried. On sites-only inputs vep-rs leads fastVEP by 4.68× to 9.43× on these two datasets, but with the genotype matrix retained the gap widens to 65.0× to 145×. Retaining 2,504 to 3,202 samples inflates vep-rs by 1.09× to 1.36× and fastVEP by 10.6× to 26.4×. vep-rs's reader cuts each line after its eight fixed columns before parsing when the output format does not re-emit the record, as the default and tab formats do not, so the sample columns are read from the file but never split or copied, while fastVEP splits every column and copies each sample column into an owned string, a per-sample allocation that scales with the sample count.
